# A Chemokine-Fusion Vaccine Targeting Immature Dendritic Cells Elicits Elevated Antibody Responses to Malaria Sporozoites in Infant Macaques

**DOI:** 10.1101/530840

**Authors:** Kun Luo, James T. Gordy, Fidel Zavala, Richard B. Markham

## Abstract

Infants and young children are the groups at greatest risk for severe disease resulting from Plasmodium falciparum infection. We previously demonstrated in mice that a protein vaccine composed of the chemokine macrophage inflammatory protein 3α genetically fused to the minimally truncated circumsporozoite protein of P. falciparum (MCSP) elicits high concentrations of specific antibody and significant reduction of liver sporozoite load in a mouse model system. In the current study, a squalene based adjuvant (AddaVax, InvivoGen, San Diego, Ca) equivalent to the clinically approved MF59 (Seqiris, Maidenhead, UK) elicited greater antibody responses in mice than the previously employed adjuvant polyinosinic:polycytidylic acid, ((poly(I:C), InvivoGen, San Diego, Ca) and the clinically approved Aluminum hydroxide gel (Alum, Invivogen, San Diego, Ca) adjuvant. Use of the AddaVax adjuvant also expanded the range of IgG subtypes elicited by mouse vaccination. Sera passively transferred into mice from MCSP/AddaVax immunized one and six month old macaques significantly reduced liver sporozoite load upon sporozoite challenge. Protective antibody concentrations attained by passive transfer in the mice were equivalent to those observed in infant macaques 18 weeks after the final immunization. The efficacy of this vaccine in a relevant non-human primate model indicates its potential usefulness for the analogous high risk human population.

---

Infants and young children are the groups at greatest risk for severe disease resulting from *Plasmodium falciparum* infection. We previously demonstrated in mice that a protein vaccine composed of the chemokine macrophage inflammatory protein 3α genetically fused to the minimally truncated circumsporozoite protein of *P. falciparum* (MCSP) elicits high concentrations of specific antibody and significant reduction of liver sporozoite load in a mouse model system. In the current study, a squalene based adjuvant (AddaVax, InvivoGen, San Diego, Ca) equivalent to the clinically approved MF59 (Seqiris, Maidenhead, UK) elicited greater antibody responses in mice than the previously employed adjuvant polyinosinic:polycytidylic acid, ((poly(I:C), InvivoGen, San Diego, Ca) and the clinically approved Aluminum hydroxide gel (Alum, Invivogen, San Diego, Ca) adjuvant. Use of the AddaVax adjuvant also expanded the range of IgG subtypes elicited by mouse vaccination. Sera passively transferred into mice from MCSP/AddaVax immunized one and six month old macaques significantly reduced liver sporozoite load upon sporozoite challenge. Protective antibody concentrations attained by passive transfer in the mice were equivalent to those observed in infant macaques 18 weeks after the final immunization. The efficacy of this vaccine in a relevant non-human primate model indicates its potential usefulness for the analogous high risk human population.

The World Health Organization reported in 2019 that estimated worldwide malaria incidence continued along a path of annual increases from a nadir of 212 million cases in 2015 to 228 million cases in 2018 ^1^. This ongoing expansion of the epidemic and emerging evidence of increased resistance to antimalarial drugs ^2,3^ indicate the continued need for cost-effective strategies to prevent malaria infection. The development of an effective malaria vaccine would address this need, and studies in mouse model systems provide clear evidence that sufficient concentrations of specific antibody can protect against malaria infection ^4–10^. However, candidate vaccines tested in human populations have failed to achieve and sustain the levels of protection expected from an effective vaccine ^11^. The inadequacy of the observed protection has been particularly apparent in the infants and young children who are at greatest risk of severe malaria infection ^11,12^.

Reports that prolonging the interval between primary and booster immunizations, as well as reducing the vaccine dose used in secondary immunizations, may provide a strategy for extending protection with the RTS,S malaria vaccine that has undergone advanced testing in clinical trials ^11,13,14^. However, these strategies have not addressed the finding that young infants respond poorly to this vaccine ^11,12,15^.

We have previously reported on the protective efficacy in a mouse challenge model of a vaccine platform that fused the chemokine macrophage inflammatory protein 3α (MIP3α), also known as Chemokine (C-C Motif) Ligand 20 (CCL20), to a slightly truncated version of the circumsporozoite protein of the *Plasmodium falciparum* parasite (PfCSP). When used with the experimental adjuvant polyinosine-polycytidylic acid (poly (I:C)) this vaccine construct induced a response that reduced liver sporozoite load by 30-fold ^16^. The 30-fold reduction in sporozoite load following challenge with transgenic *Plasmodium berghei* sporozoites expressing a region of the *P. falciparum* circumsporozoite protein was sustained for 23 weeks following the final immunization, the latest time point tested ^16^. Protection sustained over that period of time post final immunization had not previously been reported in malaria mouse challenge models.

The MIP3α fusion component of this vaccine construct has two functions: 1) To target the vaccine antigen to the C-C Motif Chemokine Receptor 6 (CCR6) protein present on the surface of the immature dendritic cells (iDC) that initiate the adaptive immune response ^17,18^ and 2) To attract immune cells to the site of immunization ^16,19^. Previous studies demonstrated marked enhancement of the antibody response to this vaccine construct compared to that observed with vaccine constructs not employing the chemokine component ^16,20,21^

In the current study we have compared in the mouse model system the immune responses and protection observed using the previously employed poly(I:C) to the research grade formulations of two clinically approved adjuvants: a squalene based adjuvant (AddaVax, InvivoGen, San Diego, Ca) equivalent to the clinically approved MF59 (Seqiris, Maidenhead, UK) and the clinically approved Aluminium hydroxide gel (Alum, Invivogen, San Diego, Ca) adjuvant. These studies indicate that the clinically approved adjuvants elicited more profound antibody responses than poly (I:C), that the addition of the MIP3α component used with the AddaVax adjuvant resulted in mice in a tripling of the maximum achieved antibody concentrations, and that the high concentrations of antibody elicited to the CSP component were not associated with any detectable antibody response to the host-derived chemokine component of the fusion vaccine construct. Further, use of the AddaVax adjuvant elicited IgG isotype responses not elicited by the Alum adjuvant. We then examined the immunogenicity of this vaccine construct used with the AddaVax adjuvant in infant and juvenile macaques. We found that this vaccine elicits antibody responses in infant macaques, which, even in this limited study with small numbers of animals, are significantly greater than those observed in older juvenile macaques. Further, sera or immunoglobulin from immunized macaques passively transferred to mice demonstrated the ability of macaque antibody concentrations persisting for 18 weeks after the final immunization to significantly reduce mouse liver sporozoite load following intravenous challenge with large sporozoite inocula.

## Materials and Methods

### Animals

Six- to eight-week-old female C57BL/6 (H-2b) mice were purchased from Charles River Laboratory (Charles River Laboratories, Inc, Wilmington, MA) and maintained in a pathogen-free micro-isolation facility in accordance with the National Institutes of Health guidelines for the humane use of laboratory animals. One and six month old Rhesus macaques were housed at the Johns Hopkins University Research Farm. All experimental procedures involving mice and monkeys were approved by the Institutional Animal Care and Use Committee of the Johns Hopkins University and determined to be in accordance with the guidelines outlined in the Animal Welfare Act, federal regulations, and the Guide for the Care and Use of Laboratory Animals.

### Construction, expression, production and purification of recombinant proteins

Recombinant *P. falciparum* CSP (3D7) and CSP fused with human MIP-3α proteins were prepared as previously described ^16^. The codon-optimized CSP DNA sequence containing deletions of the N-terminal 20 aa signaling sequence and the C-terminal 23 aa anchor region fused with human MIP-3a was used to express MIP-3α-CSP chimeric protein (MCSP). Twenty-two NANP repeats were included in the truncated CSP. CSP and MCSP sequences were cloned into pET-47b (Novagen Inc., Madison, WI) fused with 6XHis. The proteins were expressed in BL21 DE3 competent cells (New England Biolabs, Ipswich, MA). Protein purification was undertaken using nickel-affinity chromatography (Qiagen, Valencia, CA, USA) and endotoxin was removed by two-phase extraction with Triton X-114 ^22^. Protein concentration was measured by Bradford assay (BioRad, Hercules, CA). Endotoxin concentration was determined using the ToxinSensorTM Chromogenic LAL Endotoxin Assay Kit (Genscript, NJ). All protein used for immunization had final endotoxin levels below 10 EU/ml. Polyacrylamide gel electrophoresis of the final products are shown in Figure S1.

### Animal immunization

C57BL/6 mice were immunized with 20 μg (50 μl) purified MCSP in PBS with the adjuvants polyinosinic-polycytitdlic acid (Poly (I:C)), Aluminium hydroxide gel (Alum), or the squalene-based oil-in-water nano-emulsion (AddaVax,). All adjuvants were obtained from InvivoGen, San Diego, CA. For Poly (I:C) or Alum formulation, 50 μg adjuvant was mixed with 50 μl antigen at the specified doses, and the final volume ratio was 1:1. For AddaVax use, the emulsion, as supplied by InvivoGen was mixed with antigen at volumes equal to those of the antigen at the specified dose. The mixture was administered intramuscularly in both anterior tibialis muscles in a total volume of 50 μl per leg. Mice were immunized twice at a 3 week interval. Rhesus macaques were immunized with 50 μg or 250 μg purified MCSP in 250 μl PBS. The antigen was mixed with same volume of AddaVax and administered intramuscularly in both leg muscles in a total volume of 250 μl per leg. Macaques were immunized according to schedules described in the Results section.

### Antibody concentration and titer by ELISA

Humoral immune responses to the immunodominant B cell epitope of the relevant CSP protein were measured using a previously described method ^20^. Briefly, ELISA plates were coated with 2 μg of recombinant PfCSP overnight at room temperature (RT). Plates were blocked with 2% BSA for 30 min at RT and then sera at a 1:5000 dilution was added and incubated for 2 h. After washing six times, peroxidase labeled goat anti-mouse IgG or goat anti-human IgG antibodies (Santa Cruz Biotechnology Inc, Dallas, TX) were added at a dilution of 1:4000 and incubated at RT for 1 h. After washing six times, ABTS Peroxidase substrate (KPL, Gaithersburg, MD) was added for development and incubated for 1 h. Optical density at 405 nm (OD_405_) for each sample was collected using the Synergy HT (BioTek Instruments, Inc, Winooski, VT). A similar procedure was employed to determine ELISA titers, in which serially diluted serum samples were employed instead of the single 1:5000 dilution. The endpoint ELISA titers are reported in two different ways. Consistent with previous studies from our laboratory the titer is reported as the highest dilution of serum at which the average absorbance was twice the value obtained using pre-immunization serum. To compare our titer results in one month old macaques to those obtained by other investigators using an RTS,S combined with thrombospondin-related anonymous protein (TRAP) and formulated with AS02A (RTS,S+TRAP/AS02_A_) vaccine in the same age group, we also examined our titer results using as the endpoint the last dilution of sera that yielded an OD_405_ >1.00. For determination of antibody responses to MIP3α, recombinant purified mouse MIP3α (R&D Systems, Minneapolis, MN) was used (2 μg/well) to coat ELISA plates, as described for recombinant PfCSP.

### Parasites for challenge

Transgenic *P. berghei* sporozoites ^23^, carrying only the central repeat region of the *P. falciparum* CSP, were used for challenge. Sporozoites were obtained by hand dissection of salivary glands of *Anopheles stephensi* mosquitoes maintained in the Johns Hopkins Malaria Research Institute insectary. The isolated sporozoites were suspended in HBSS medium containing 1% normal mouse serum. Challenges to evaluate vaccine effect on hepatic parasite load were accomplished by tail vein injection of 5 x 10^3^ sporozoites.

### Real-time PCR for liver stage parasites

Approximately 40 hours after sporozoite challenge, livers of challenged mice were harvested and homogenized, and RNA was extracted. After reverse transcription (Applied Biosystems, Foster city, CA), Real-time PCR was used for the detection and quantification of the liver stage of Plasmodium parasites. Two pairs of specific primers were designed to amplify the parasite 18s rRNA sequence. The forward primer was 5’-TGGGAGATTGGTTTTGACGTTTATGT-3’ and the reverse primer was 5’-AAGCATTAAATAAAGCGAATACATCCTTAC-3’. Values were normalized against measurements of mouse actin mRNA in the same samples. The primers used for mouse actin were as follows: 5’-GTCCCTCACCCTCCCAAAAG-3’ (forward) and 5’-GCTGCCTCAACACCTCAACCC-3’ (reverse). The reactions were performed in a final volume of 20 μl using SYBR green PCR Master Mix (2X) from Applied Biosystems (Waltham, MA) and processed with ABI StepOne Real-time PCR system (Applied Biosystems).

### Passive antibody or serum transfer

Rhesus Macaques were immunized with 50 or 250 μg MCSP and 250 μl AddaVax at the indicated frequencies and intervals (see Results Section). Four weeks after each immunization, macaque blood was collected and serum was isolated. All samples were pooled and some of the sera were used to obtain purified IgG, using protein A affinity chromatography (GE Healthcare, Marlborough, MA). IgG from pooled samples that were collected from the macaques before immunization was purified as the control for passive antibody transfer. To determine ELISA titers achieved in mice by transfer of macaque sera, 200 μl of macaque sera with a reciprocal ELISA titer of 512,000 was inoculated by tail vein injection into three C57Bl/6 mice and mice were bled by tail vein 15 and 60 minutes post tail vein inoculation. The mouse sera at both time points in all three mice had a reciprocal titer of 64,000, an 8-fold dilution of the injected macaque sera. Subsequent challenge studies in mice assumed this 8-fold dilution for evaluating protective capabilities in mice of titers achieved in the macaques. For challenge studies C57BL/6 mice were injected by tail vein with 200μl purified IgG from immunized macaques or immune sera or control IgG or sera. Purified IgG was used in those settings in which higher concentrations of antibody in volumes appropriate for IV injection were required to attain the desired high titer in the mouse. Thirty minutes following injection of antibody or sera, mice were inoculated via tail vein with 5 x 10^3^ transgenic *P. berghei* sporozoites. Liver stage sporozoite copy number was measured 48h following infection.

### Statistical analysis

Linear regression analysis was used to compare the slopes of antibody decline in sera from recipients of different adjuvants. A mixed effects model analysis was used to compare OD_405_ antibody concentrations achieved with different adjuvants at individual time points after immunization. 2-way ANOVA with Tukey’s multiple comparisons test was used to analyze antibody subtypes. Welch’s ANOVA test was used to compare RNA copy number in livers of recipients of vaccine plus different adjuvants. Student’s unpaired t test was used to compare antibody responses with and without the MIP3α component and was also used to compare mouse liver sporozoite copy numbers following challenge and passive transfer of antibodies or serum from immunized macaques. Prism 8 (GraphPad Software, Inc., San Diego, CA) or STATA (version 11.2) (StataCorp LLC, College Station, TX) statistical analysis and graphing software was used for all studies.

## Results

### Evaluating the efficacy of different adjuvants in enhancing the anti-CSP humoral immune response

While vaccine adjuvant development is an active research area, only alum and MF59 (Seqiris, Maidenhead, UK), the clinical equivalent of AddaVax, have been approved by the US FDA for use in commonly employed parenterally administered vaccines. We first compared in C57Bl/6 mice the CSP-specific humoral immune responses elicited by the MCSP vaccine in combination with these two adjuvants and the previously studied poly (I:C) ^16^. The results (Figure 1) indicate that the most profound response is elicited by AddaVax, with a mean OD_405_ value significantly greater than that observed with poly (I:C) at all time points after the priming immunization (p<0.001 for all points). However, alum and AddaVax differed significantly only at 13 and 16 weeks after primary immunization, although those differences approached significance (0.1>p>0.05) at weeks 19-25.

**Fig.1.**
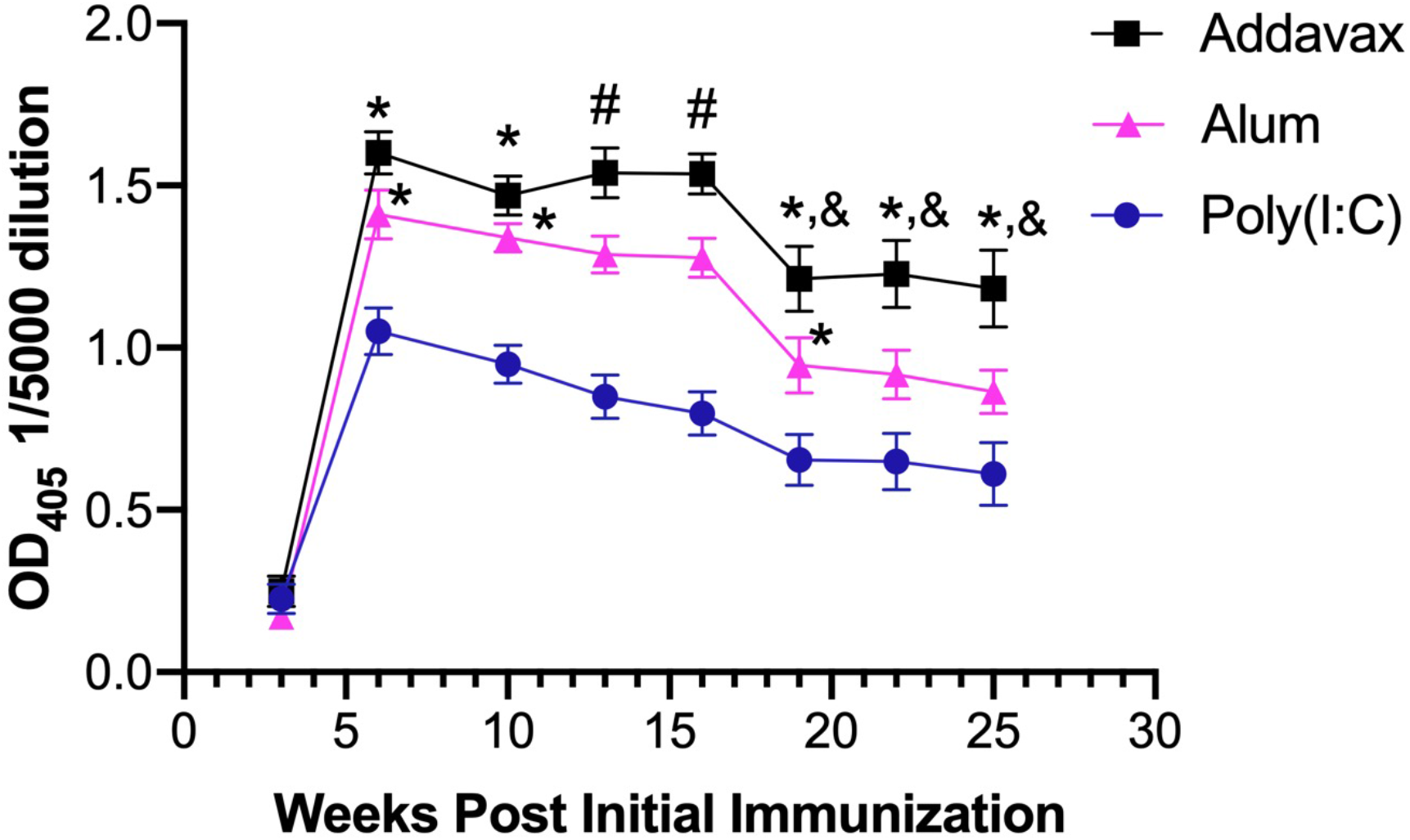
Longitudinal analysis of responses to MCSP vaccine used with different adjuvants. Groups of 15 C57BL/6 mice were immunized twice with 20 μg MCSP plus the indicated adjuvants at a three week interval. Mice were bled at the indicated time points to determine specific antibody concentrations. Values shown represent absorbance in an ELISA assay at OD_405_ nm after a 1/5000 serum dilution. By linear regression analysis of antibody response curves, the slopes of decline in antibody concentrations from the six week time point for all of the adjuvants do not differ significantly (p>0.05) from each other, although the difference in slopes between AddaVax and poly (I:C) approach significance (p=.0538). Using a Mixed Effects Model analysis, comparisons of OD_405_ levels at individual time points indicated levels achieved using AddaVax are significantly greater than those achieved with alum or poly (I:C) as indicated (*p<0.05 to poly(I:C); #p<0.05 for all group comparisons;&0.05<p<0.1 comparing Alum and Adddavax). Error bars represent Mean +SEM.

We next examined the impact of the different adjuvants on the IgG subtype of the elicited antibody response (Figure 2). Three weeks after the second immunization, elicited IgG1 responses were statistically equivalent between Alum and AddaVax, which for both of those adjuvants were significantly greater than the responses obtained using poly(I:C). The most striking difference among the adjuvants was the markedly reduced IgG2b and IgG2c responses observed with use of the alum adjuvant, which was significantly reduced compared to responses elicited with AddaVax or poly(I:C). No differences among the adjuvants was observed with the elicited IgG3 responses.

**Fig. 2.**
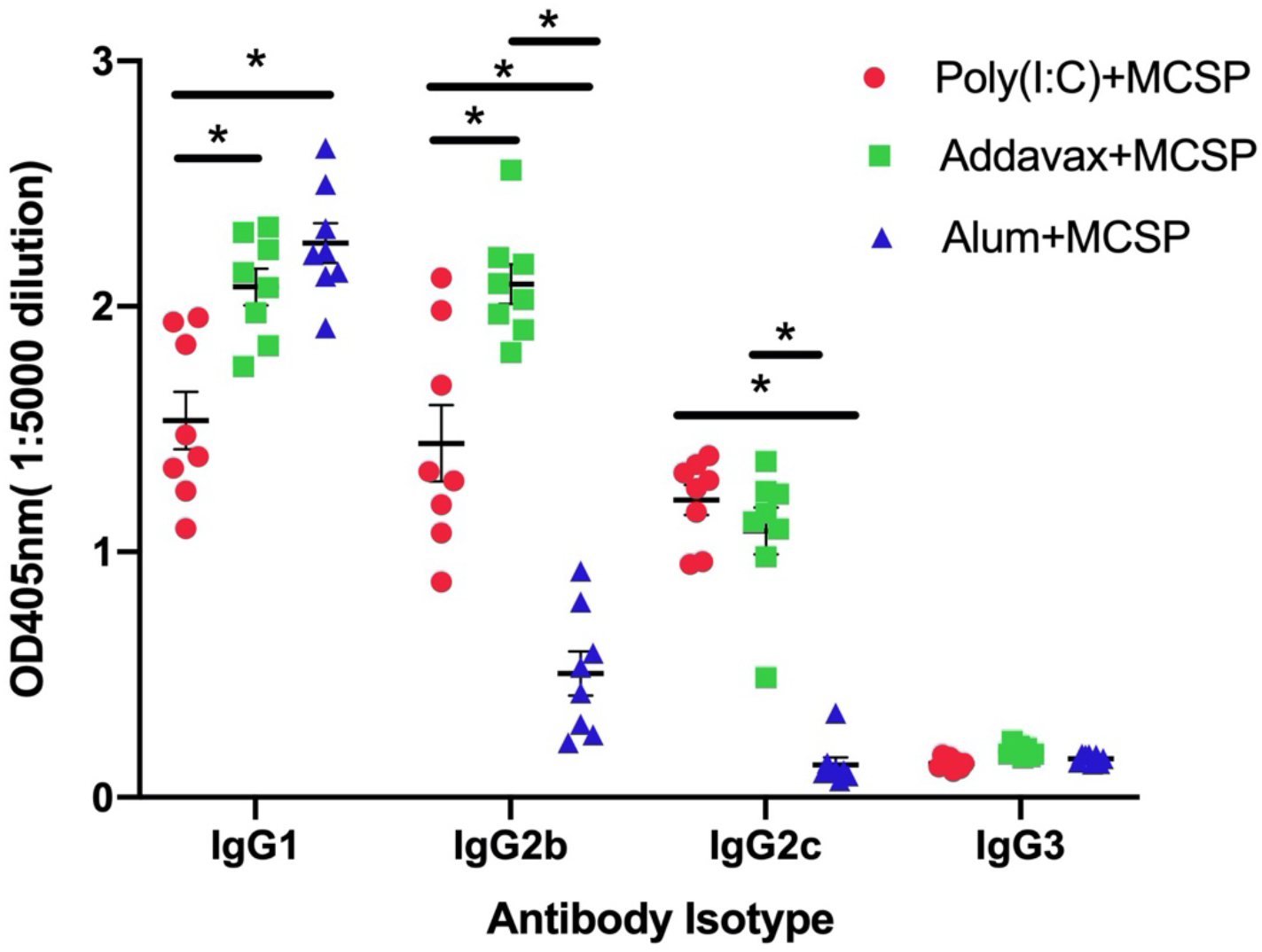
Generation of anti-CSP antibody of different isotypes after immunization with different adjuvants. C57BL/6 mice were immunized twice with 20 μg MCSP plus different adjuvants at a three week interval. Mouse sera were obtained 3 weeks after the last immunization for ELISA, performed as in Fig. 1, except that isotype specific secondary antibodies were used to determine concentrations of IgG1, IgG2b, IgG2c and IgG3. Data analyzed by 2-way ANOVA (*p<0.02). Error bars represent Mean +SEM.

### Efficacy of MIP3α-CSP vaccine combined with different adjuvants in protecting mice from mosquito bite challenge

To characterize at 22 weeks post-the final of two immunizations the protective efficacy of the different combinations of adjuvant and vaccine, immunized or non-immune C57Bl6 mice were challenged by exposure for 10 minutes to 5 *Anopheles stephensi* mosquitoes infected with transgenic *P. berghei* sporozoites. Sporozoite copy numbers determined by qRT-PCR indicated no statistically significant difference among the adjuvant groups in their impact on reduction of liver sporozoite load (p>0.18, Figure 3). However, all groups receiving vaccine plus adjuvant had highly significant reductions of at least 78% in the liver sporozoite copy number compared to unimmunized mice (p<0.01). Based on the possibility that distinct functional capabilities of different IgG isotypes in might be more important in the human setting than is the case in mice, discussed below, and also based on the significantly higher concentrations of antibody observed with use of AddaVax, a decision was made to use that adjuvant in the subsequent macaque studies.

**Fig. 3.**
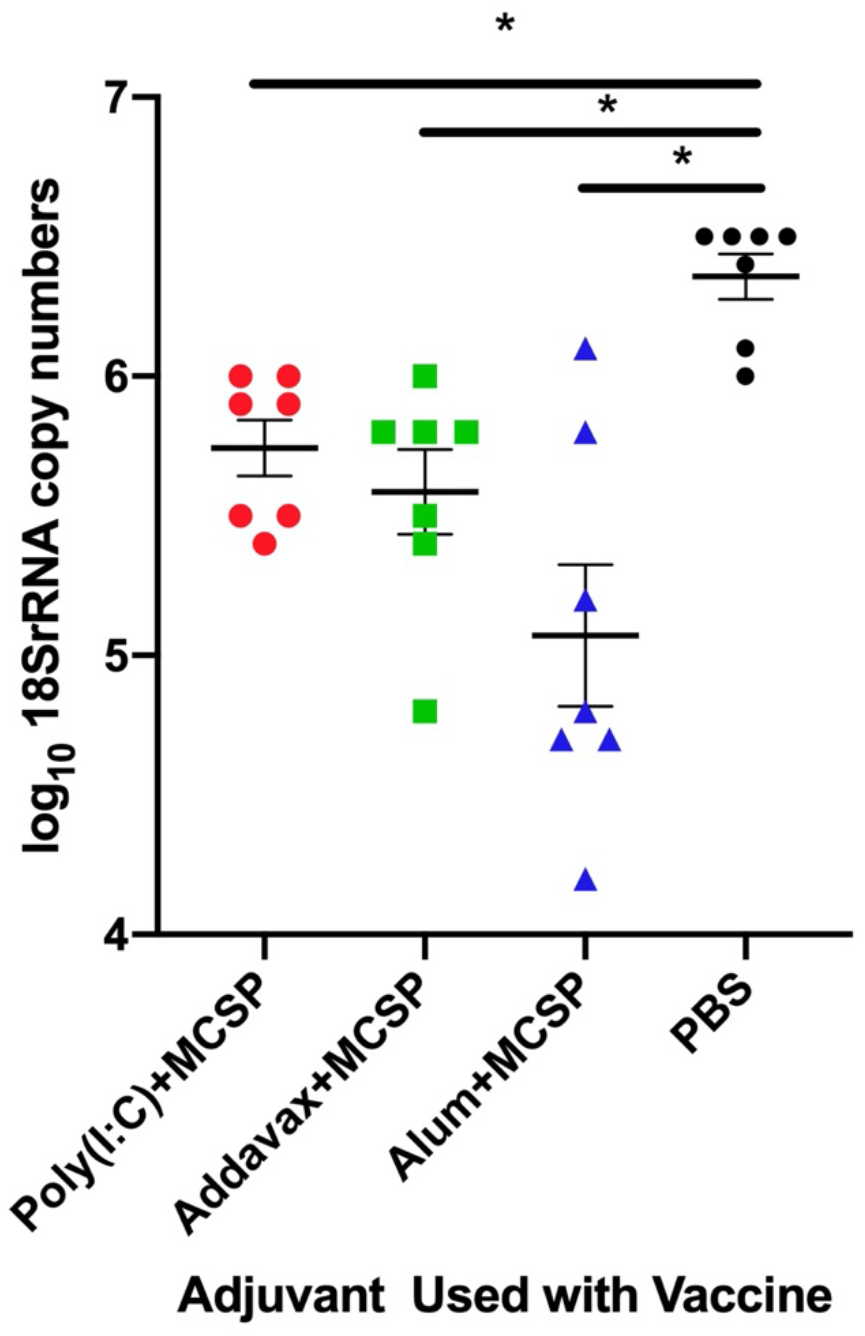
Reduction in liver sporozoite copy number in C57Bl/6 mice immunized with different adjuvants following challenge by bites of mosquitoes carrying transgenic *P. berghei*. C57BL/6 mice were immunized twice with 20 μg MCSP plus different adjuvants at a three week interval. Twenty-two weeks after the final immunization, mice were exposed for 10 minutes to 5 mosquitos infected with transgenic *P. berghei* sporozoites. Parasite-specific rRNA levels in the liver were determined by quantitative RT-PCR on samples obtained 48 hours post challenge. All qPCR results were normalized against the expression of mouse β-actin. Sporozoite copy numbers did not differ significantly among immunized mice (p>0.5) and all immunized groups differed significantly from control mice (p<0.001).

### Confirmation of benefit of the fusion construct when combined with AddaVax adjuvant

The consistently higher antibody concentrations achieved with AddaVax and its ability to elicit a broader subclass response led to its selection for use in subsequent studies. Before proceeding with further studies, we sought to confirm that with use of the AddaVax adjuvant, we still observed the benefit in immunogenicity and protection with the fusion vaccine compared to use of adjuvant and fCSP vaccine without the fused MIP3α component. For this purpose, antibody titers were determined two weeks after the last of two immunizations using the AddaVax adjuvant with either the CSP or MCSP vaccine construct. CSP-specific antibody concentrations were approximately three-fold higher in recipients of the MCSP vaccine (Figure 4a, p=0.01). Following intravenous challenge with 5 x 10^3^ sporozoites three weeks after the final immunization, there was an approximately one log greater reduction in the liver sporozoite copy number in the MCSP immunized group compared to the CSP immunized group (Figure 4b, p=0.02).

**Figure 4.**
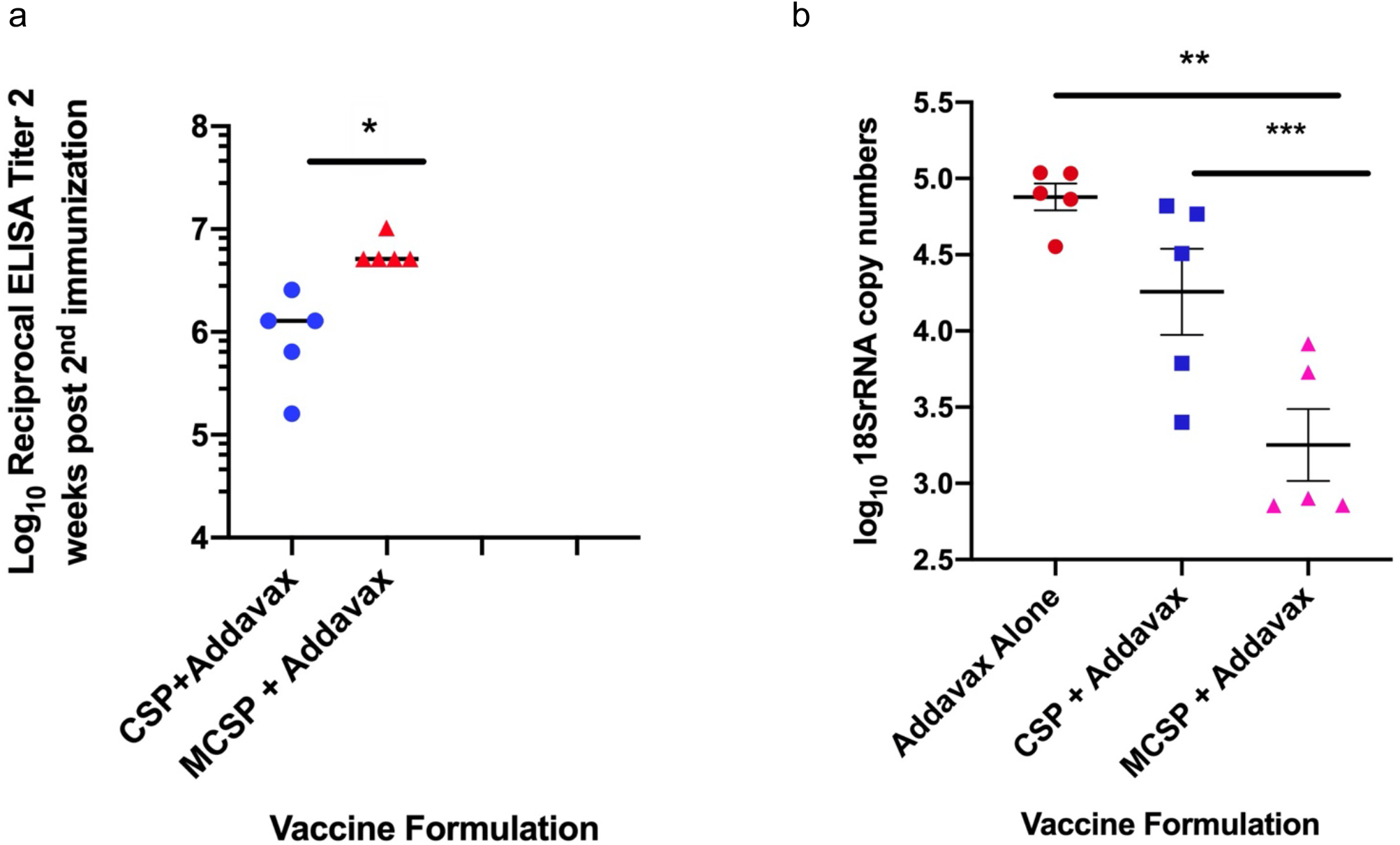
Effect of fusion of MIP3α to CSP on antibody response and vaccine-mediated reduction in liver sporozoite load when used in combination with AddaVax adjuvant. **a.** C57BL/6 mice were immunized twice at a three week interval with 20ug of the indicated antigen adjuvanted with AddaVax (1:1/v:v). The antibody concentration was determined 2 weeks after the second immunization. The endpoint ELISA titer is reported as the highest dilution of serum at which the average absorbance was twice the value obtained using the average pre-immunization sera. **b.** C57BL/6 mice were immunized with 20ug of the indicated antigen adjuvanted with AddaVax (1:1/v:v). 3 weeks after the second immunization, mice were challenged with 2000 transgenic sporozoites. Parasite 18S rRNA copy number was determined by qPCR. * p<0.004, ** p=0.005, *** p=0.02. Error bars represent Mean +SEM.

### Determination of whether MCSP vaccine elicits a humoral immune response directed at the MIP3α vaccine component

Because of the apparent immunogenic potential of this vaccine formulation, we examined whether it might elicit an antibody response directed at the fused MIP3α vaccine component. The vaccine used in the previous studies employed human MIP3α in the vaccine, which binds murine CCR6. While human and murine MIP3α both bind to murine CCR6 ^24^, the MIP3α from these two species are only 70% homologous at the protein level ^25^. Previous studies have demonstrated that mice failed to generate an antibody response to human MIP3α with use of the vaccine plus poly(I:C) adjuvant ^16^, but given the enhanced immunogenicity observed with the AddaVax adjuvant we sought to confirm that finding, using a vaccine construct in mice containing the homologous mouse MIP3α. A vaccine construct employing murine MIP3α was developed and its ability to elicit anti-CSP and anti-mouse MIP3α antibodies was studied two weeks after the last of two immunizations, the point of maximal humoral immune response. This analysis revealed that at that time point the mean OD_405_ reading for anti-MIP3α antibody concentration at a serum dilution of 1/200 was 0.068, while that for the pre-immune sera at that dilution was 0.060. The same group of mice had a mean OD_405_ reading of 0.144 for anti-CSP antibody after being diluted 640,000 fold (Figure 5). Despite its ability to elicit high concentrations of antibody to the malaria CSP antigen, this vaccine construct does not elicit an antibody response to the host-derived chemokine component.

**Figure 5.**
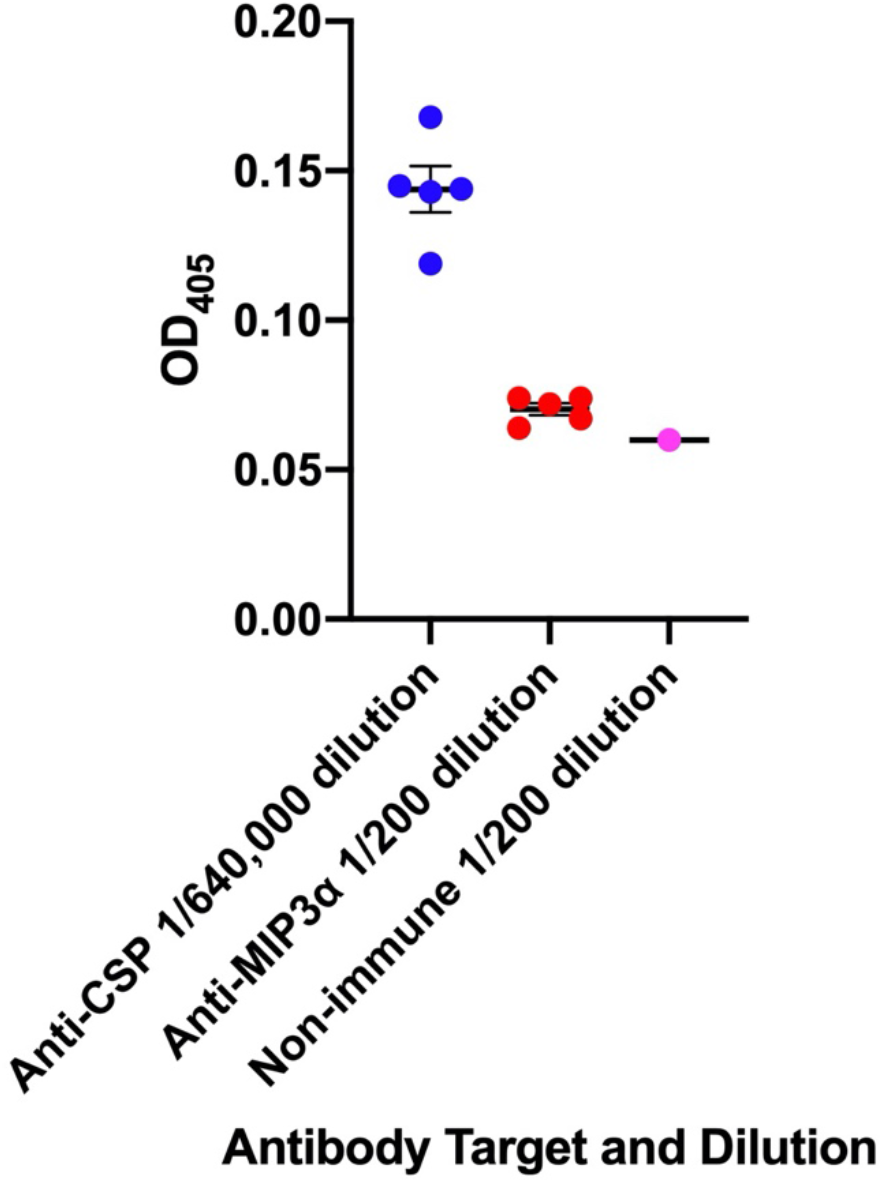
Antibody Response to MIP3α and CSP Vaccine Components. Five C57BL/6 mice were immunized at a three week interval with 20ug of the MCSP vaccine adjuvanted with AddaVax (1:1/v:v). The antibody concentration evaluated by OD_405_ readings was determined 2 weeks after the second immunization. The endpoint ELISA titer for the anti-CSP response is reported as the highest dilution of serum at which the average absorbance was twice the value obtained using the mean absorbance of preimmunization sera. Open circles represent the OD_405_ readings from a pool of pre-immune serum at the 1/200 dilution. Error bars represent Mean +SEM.

### Immunogenicity of MCSP vaccine and AddaVax adjuvant in one and six month old macaques

One issue in vaccine development is whether responses in mice can be recapitulated in the clinical setting. Non-human primates, particularly Old World primates such as rhesus macaques, have immune systems that are most homologous to humans ^26–30^. To address the immunogenicity of this vaccine construct in a setting relevant to the groups at high risk from malaria, we initiated an immunization protocol in infant, one-month-old, and juvenile, six-month-old, macaques, using two dosing regimens. In each of the age groups two macaques received a priming 50 μg dose of the MCSP vaccine construct and 250 μl of the AddaVax adjuvant followed by two subsequent immunizations with the same dosing at monthly intervals. The two other macaques in each age group received a priming immunization with 250 μg of vaccine and 250 μl of the AddaVax adjuvant followed one month later by a second immunization using the same vaccine and adjuvant dose. Figure 6 displays the antibody responses, expressed as reciprocal ELISA titers, maintained over 25 weeks post the initial immunization. For 18 weeks after the final immunization, all of the infant macaques, independent of the immunization regimen, maintained antibody titers that, when replicated by passive transfer in C57Bl/6 mice, significantly reduced liver sporozoite load (see passive protection studies below). Juvenile macaques maintained antibody titers associated with reduced liver sporozoite loads in passively immunized mice for between 10 and 18 weeks.

**Figure 6.**
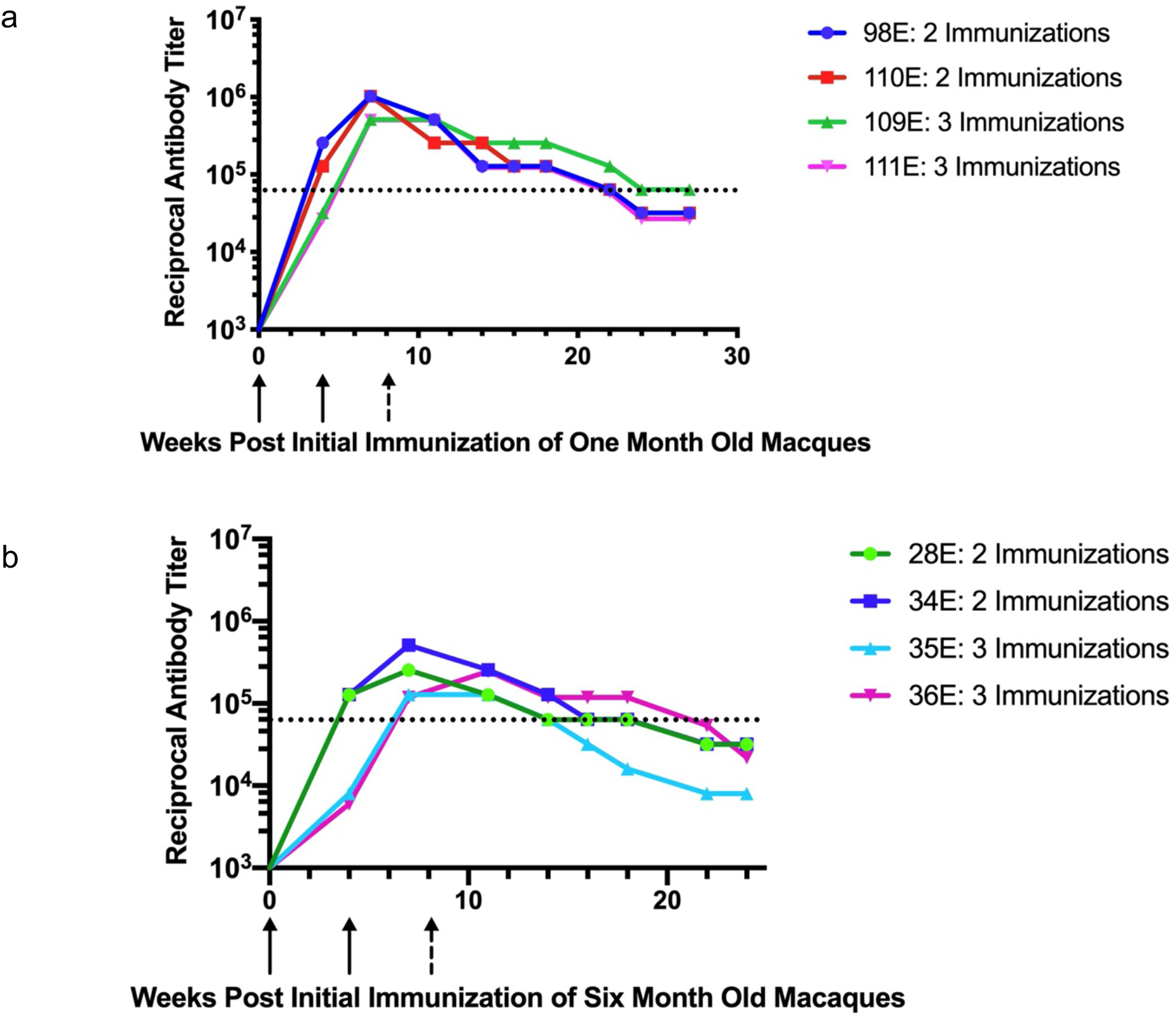
Response of One and Six Month Old Rhesus Macaques to Different MCSP + AddaVax Immunization Regimens. a) One month old macaques 98E and 110E received two immunizations with 250 μg MCSP + 250 μl AddaVax, indicated by solid arrows. One month old macaques 109E and 111E received three immunizations with 50 μg MCSP + 250 μl AddaVax (solid arrows for first two immunizations and broken arrow for the third immunization). b) Six month old macaques received similar regimens: 28E and 34E received two immunizations with 250 μg MCSP + 250 μl AddaVax, indicated by solid arrows. Macaques 35E and 36E received three immunizations with 50 μg MCSP + 250 μl AddaVax (solid arrows for first two immunizations and broken arrow for the third immunization). For both figures, the dotted black line indicates reciprocal antibody titer associated with significant reduction in liver sporozoite load in C57Bl/6 mice passively administered sera or immunoglobulin from immunized macaques (see Figure 8). Positive titer = 2x the pre-immunization control.

### Comparison of antibody responses in one and six month-old macaques

A recent study of responses in humans to a candidate HIV-1 vaccine found that the vaccine adjuvanted with AddaVax elicited higher responses in human infants than those observed in adults ^31^. In the current study, the peak antibody concentrations obtained three weeks after the second immunization, evidenced no significant dose-dependent differences in the observed responses across both age groups (p=0.13), suggesting that an immunogenicity threshold had been reached with the 50 μg vaccine dose. We therefore compared the responses between the age groups, combining recipients of both immunizing doses for this analysis (Figure 7). Even with the small sample size of this immunogenicity study, the maximum antibody response obtained three weeks after the second immunization was significantly higher in macaques initially immunized at one month of age compared to those initially immunized at six months of age (p=0.03).

**Figure 7.**
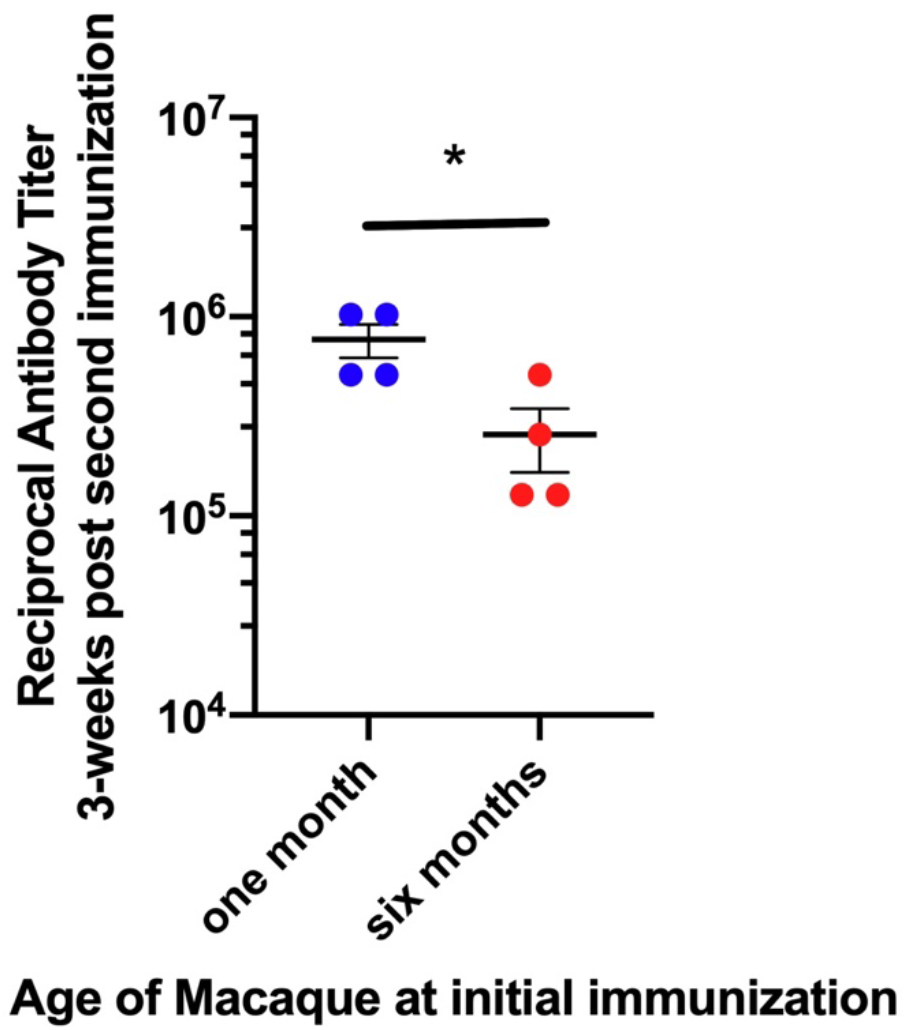
Macaques initially immunized at one month of age generate greater antibody responses. Mice were immunized as in Figure 6 and were evaluated by age at initial immunization independent of immunizing regimens. At 7 weeks after the initiation immunization, the point of greatest response, the antibody concentrations were significantly higher in the macaques initially immunized at one month of age compared to those initially immunized at 6 months of age (p=0.03).

### Passive protection by immune macaque sera in a mouse challenge model

To evaluate the protective capability of the different antibody concentrations achieved in the Rhesus macaques, immune macaque sera or purified immune macaque IgG was passively transferred to naïve C57Bl/6 mice that were challenged via intravenous inoculation with transgenic *P. berghei* sporozoites expressing *P. falciparum* CSP. This challenge model was used because there are no macaque pre-erythrocytic challenge models for *P. falciparum*. 200 μl IgG or sera were administered in concentrations to achieve in the mice a titer equivalent to the titers achieved in macaques at different time points after immunization. As indicated, this intravenous injection of 200 μl of macaque IgG or sera resulted in an 8-fold dilution in mouse sera of the concentration of the inoculated macaque sera (see Methods). Control mice received pre-immune macaque sera or IgG in volumes equivalent to that received by recipients of immune sera or IgG. Mice with reciprocal anti-CSP ELISA titers as low as 64,000 still had significant reduction in their liver sporozoite load (Figure 8).

**Figure 8.**
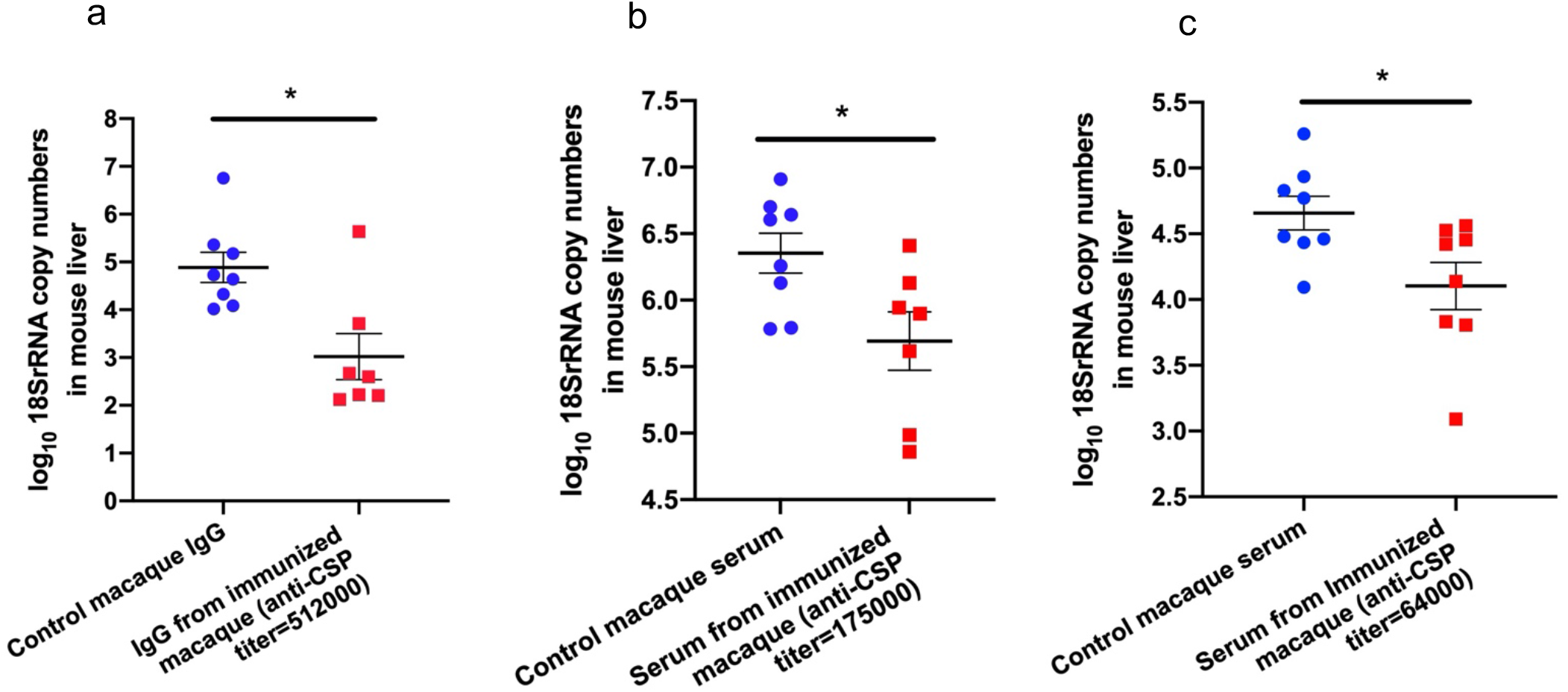
Protection of C57Bl/6 mouse recipients of sera or immunoglobulin from immunized infant or juvenile macaques against intravenous challenge with transgenic *P. berghei* expressing *P. falciparum* CSP. C57BL/6 mice were injected i.v. with 200 μl IgG from MCSP/AddaVax immunized macaques (a) or sera from MCSP/AddaVax immunized macaques (b,c). Groups of control mice were injected with IgG purified from sera (a) obtained from macaques prior to immunization or directly with the pre-immunization sera (b). IgG was used in (a) to reduce the volume injected into the mice to attain the 512,000 reciprocal endpoint dilution anti-CSP titer. Indicated titers are those calculated to have been attained in the mice after injection of IgG or serum of differing concentrations (see Methods). 30 minutes following injection of IgG or sera, mice were infected i.v. with 5 × 10^3^ transgenic *P. berghei* sporozoites expressing Pf CSP. Parasite-specific 18S rRNA levels in the liver were determined by qPCR on samples obtained 48h post challenge. Error bars represent Mean +SEM. *p<0.02 by unpaired Student’s t-test.

### Comparison in one month old macaques of responses to MCSP vaccine to previously reported responses to RTS,S+TRAP/AS02_A_

Any malaria vaccine development should involve comparisons with the RTS,S vaccine currently undergoing continuing clinical trials. Unfortunately, direct comparisons in pre-clinical models cannot be undertaken due to the refusal of the manufacturer of the RTS,S vaccine to make it available for comparison studies. However, a study by Walsh *et al*.^32^ involving one month old macaques receiving an RTS,S+TRAP/AS02_A_ vaccine formulation employing immunization protocols similar to ours has been published and provides some basis for comparison of the two vaccine formulations in this pre-clinical setting. In those studies the investigators employed a different endpoint criterion (OD_405_>1.0) for determining ELISA endpoints. For purposes of comparison we used the same endpoint to express the data shown in Figure 6 to generate comparable endpoints and to identify, using that approach, the titer associated with protection in the mice (Figure 9a). The data used from Walsh *et al*. is extrapolated from figures provided in that publication, which did not present standard deviation or standard error information. Therefore, no statistical comparison can be made with the MCSP vaccine. Nevertheless, the data presented in Figure 9b indicate that antibody titers achieved with the RTS,S+TRAP/AS02_A_ vaccine generally fell below those associated with protection in the mouse passive transfer assays.

**Figure 9.**
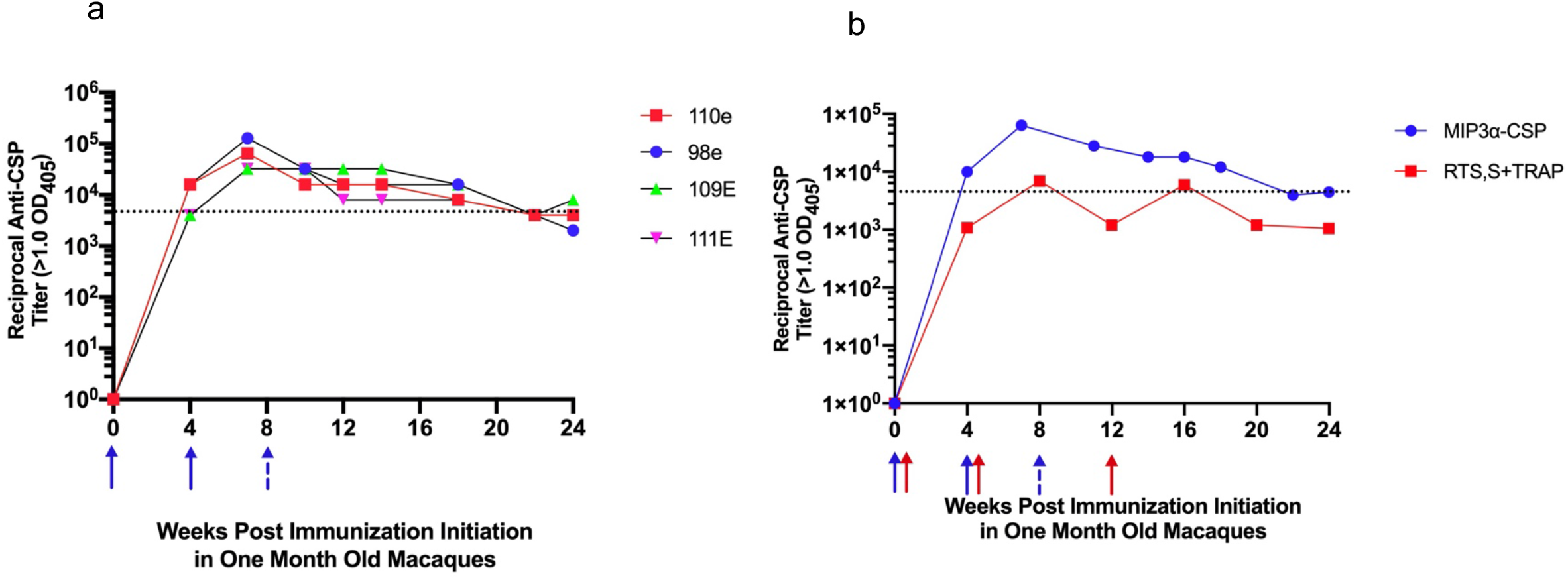
Responses in One Month Old Rhesus Macaques to MCSP + AddaVax Immunization Regimens Compared with Previously Reported Results with RTS,S+TRAP/AS02_A_ Vaccine Using Titers with OD_405_>1.00 as Dilution Endpoint. One month old macaques 98E and 110E received two immunizations with 250 μg MCSP + 250 μl AddaVax, indicated by solid arrows. One month old macaques 109E and 111E received three immunizations with 50 μg MCSP + 250 μl AddaVax (solid arrows for first two immunizations and broken arrow for the third immunization). The dotted black line indicates reciprocal antibody titer associated with significant reduction in liver sporozoite load in C57Bl/6 mice passively administered sera or immunoglobulin from immunized macaques (see Figure 8). Positive titer = OD_405_>1.00. b) Longitudinally obtained mean titers of all macaques initially immunized at one month of age with MCSP +AddaVax or RTS,S+TRAP/AS02_A_ vaccines. Titers for the RTS,S+TRAP/AS02_A_ vaccine are extrapolated from a figure in Walsh *et al*.^32^.

## Discussion

In the current study, we have examined in C57Bl/6 mice and infant and juvenile Rhesus macaques the immunogenicity of a malaria vaccine platform that had previously demonstrated, using a different adjuvant, extended protection efficacy in a murine malaria challenge system. A distinctive feature of the vaccine platform is its fusion of vaccine antigen to the chemokine MIP3α, which serves as the ligand for CCR6, expressed on iDC. The current study confirmed that, with use of the AddaVax adjuvant in a mouse model system, fusion of human MIP3α to the CSP antigen enhanced the immunogenicity of the vaccine construct. Although both are able to bind to mouse CCR6, mouse and human MIP3α are only 70% homologous at the amino acid level, but previous studies had demonstrated that even with that level of difference, an immune response to the human MIP3α was not elicited ^16^. In the current study, we demonstrated that with use of the AddaVax adjuvant, vaccination with MCSP did not elicit an immune response in mice to the autologous MIP3α, while a high titer antibody response was elicited to the CSP component. Human and macaque MIP3α are 90% homologous at the amino acid level. We did not test in the current study whether an immune response was elicited in macaques to the human MIP3α used in our vaccine construct, but considering the studies in mice, it is unlikely that such a response would be generated.

The AddaVax adjuvant selected for study in the macaques was chosen based on its ability to elicit high concentrations and diverse IgG isotypes of specific antibodies in the mouse model system and the fact that its clinical homologue, MF59, has been approved as an adjuvant for clinical use with other vaccines by the United States Food and Drug Administration. This vaccine construct and adjuvant target the earliest steps in the encounter between the host immune system and the foreign antigen. Those steps include the attraction of antigen presenting cells to the site of immunization, attributable to the chemoattractant properties of both AddaVax and MIP3α ^16,33^, and the efficient targeting of antigen to the CCR6 receptor on iDC that are typically poorly represented at immunization sites, despite the use of adjuvants ^20 34–36^. Previous studies with this vaccine used with the adjuvant poly(I:C) have demonstrated the independent chemoattractant activity of each component, but also synergistic chemoattractant ability when both the MIP3α construct and the adjuvant were employed ^16,20^. In addition to being expressed on iDC, CCR6 is also expressed on memory effector CD8+ T cells, regulatory FoxP3+ CD4+ T cells, and Th17 CD4+ T cells ^37–39^. Since these are all memory cells, it is unlikely that such cells with CSP specificity would be present in muscle at the site of primary immunization. While there is a theoretical concern that CCR6^+^ regulatory T cells with CSP specificity might be attracted to the immunization site at the time of secondary immunization or in malaria-exposed individuals, we observe marked enhancement of antibody concentrations following secondary immunization in both mice and macaques.

Studies of the mechanism of action of AddaVax and the related MF59 adjuvant have shown that it also promotes the differentiation of monocytes into cells that bear the iDC phenotype and that it promotes release of chemokines such as CCL2 (MCP-1), CCL3 (MIP-1α), or CCL4 (MIP-1β) that would be expected to attract additional iDC to the vaccination site. Further, AddaVax enhanced expression of CCR7 is implicated in the migration of DC from peripheral tissues to lymph nodes ^40^. These specific immunoenhancing activities of AddaVax position it well to complement the activity of an iDC targeting vaccine.

The antibody concentrations attained using AddaVax as the adjuvant were uniformly higher than those attained with use of alum or poly (I:C) and those differences may be relevant in the clinical setting. Because clearance of sporozoites clearly requires high concentrations of antibody, the titers achieved with any successful malaria vaccine formulation will be critical. While the comparison of different adjuvants showed higher peak antibody concentrations with use of AddaVax, the slope of antibody decline was similar with all adjuvants, with a reduction of approximately 90% in antibody concentration after six months. The observed rate of decline in antibody concentration is similar to that observed both with the RTS,S vaccine in human trials ^12^ and with a candidate influenza vaccine combined with the AddaVax adjuvant tested in children ^41^. While this decline in antibody concentrations would be viewed as suboptimal for a clinical vaccine, studies with the RTS,S vaccine have indicated that extended elevation of antibody titers can be attained by alteration of the immunization schedule ^13,14,15^. If that approach to extending protection proves successful, it allows for a renewed focus simply on vaccine formulations attaining the highest concentrations of antibody. The higher titer of specific antibody generated in infant macaques by our vaccine formulation compared to that previously reported following RTS,S+TRAP/AS02_A_ vaccination of infant macaques indicates further studies for potential application of the MCSP + Addavax in that age group are warranted.

Vaccination with the AddaVax adjuvant also resulted in significantly higher concentrations of IgG2b and IgG2c in the sera of immunized mice, particularly when compared to alum, the other clinically approved adjuvant employed in our analysis. There are conflicting data on the importance of IgG isotype in resistance to malaria infection. Aucun *et al*. showed that elevated levels of human IgG2 specific for blood stage antigens were associated with resistance to malaria infection in Burkina Faso ^42^, while Aribot *et al*. found that only concentrations of IgG3 specific for blood stage antigens were associated with reduced risk of infection in Senegal ^43^. Studies with the RTS,S/AS01E vaccine in Mozambique and Ghana found protection against sporozoite mediated infection positively associated with concentrations of IgG1 and IgG3 antibodies specific for vaccine antigens ^44^. These human antibody isotypes are important in mediating antibody-dependent cellular cytotoxicity and complement dependent killing, as are the mouse IgG2b and 2c isotypes elicited by AddaVax, but not alum, in the current study. Of note, Kurtovic *et al*. ^45,46^ found that complement fixing capability of sera from immunized infant and child recipients of the RTS,S vaccine declined rapidly after immunization and this decline paralleled the decline both in IgG complement fixing isotypes and protection against severe disease. While this finding suggests further justification for the use of the AddaVax adjuvant in subsequent vaccine trials, Potocnjak *et al*. found in mouse challenge models that Fab fragments of monoclonal antibodies to a sporozoite surface antigen, which would lack complement fixing capability, could protect mice against sporozoite challenge at concentrations roughly equivalent to protective concentrations of fully intact antibodies ^47^. This apparent difference in the importance of complement fixing antibodies in malaria infection in human and murine settings may be attributable to the fact that circulating complement (CH50) levels are approximately 40-fold greater in humans than in the Balb/c or C57Bl/6 mice typically used in malaria studies ^48^. Whether the broader isotype response achieved with AddaVax provides additional protection in the clinical setting will, therefore, require further study.

As indicated, a recent study of responses in humans to a candidate HIV-1 vaccine conducted using the AddaVax adjuvant demonstrated higher responses in human infants than those observed in adults ^31^. This observation, which was shown not to be attributable to dose/body mass calculations, is consistent with our findings and suggests that AddaVax may be a particularly useful adjuvant for use in infants and very young children.

Macaques were chosen for our nonhuman primate study because their immune ontogeny more closely parallels that of humans than does that of rodents or that of the New World monkeys that have been used as non-human primate challenge models of malaria bloodstream infection ^26–30^. A study comparing the immunogenicity of two formulations of a candidate sporozoite-targeted vaccine demonstrated that different conclusions might be reached using mouse vs macaque models for analysis of CSP immunogenicity ^49^. Pre-erythrocytic stage *P. falciparum* challenge models using macaques have been unsuccessful, and efforts to develop *Aotus* monkey based challenge systems have also proved problematic. Reports from two laboratories indicated success with sporozoite transmission in this species ^50,51^. However, for sporozoite transmission to Aotus monkeys, splenectomy is required and, according to these reports, successful transmission occurs with varying frequency, achieves highly variable levels of parasitemia, has wide variation in the pre-patent period, is often accompanied by clearing of parasitemia in the non-immune host and requires use of parasites that have been obtained from relatively recent human infection for best results ^50^. No studies using this model have been published since 2009.

To evaluate the protective efficacy of the antibody concentrations attained with macaque immunization we determined the quantity and ELISA titer of passively transferred IgG or serum that in mice could duplicate a specific concentration in macaques. In some groups purified and concentrated IgG was required to reproduce in the mice the higher concentrations observed in the macaques. For these studies the endpoint titer for dilution was defined as 2x the negative control. Passively immunized mice were challenged with inocula of 5 x 10^3^ transgenic *P. berghei* sporozoites. The results indicated that a reciprocal macaque antibody titer in the mice of 6 x 10^4^ still yielded a significant reduction in liver sporozoite load. Using that criterion, antibody titers associated with significant reduction in liver sporozoite load were maintained for 18 weeks after a second immunization in the limited number of infant macaques evaluated (Figures 6 and 9a).

Because the RTS,S vaccine has been the most extensively studied in the clinical setting ^14,52–55^, alternative vaccine platforms would ideally be compared to that vaccine formulation. As stated, the vaccine constructs used in those clinical studies are not being made available for comparison studies. Using previously published data that employed an RTS,S vaccine formulation in one month old macaques immunized according to a similar protocol, we were able to provide evidence that the MCSP + AddaVax vaccine formulation generated higher concentration of antibodies than the RTS,S formulation (Figure 9b). Critically, unlike the MCSP + AddaVax immunization regimen, the RTS,S+TRAP/AS02_A_ vaccine, was able to only very transiently elicit antibody titers associated with significant liver sporozoite load reduction in our mouse challenge system (Figure 9b). This comparison is admittedly imperfect in that 1) the RTS,S vaccine also included a second antigen, TRAP, the effect of its presence in the vaccine construct being unknown and 2) the unavailability of the variation in the published RTS,S + TRAP results prevents any statistical analysis of the significance of the observed differences.

The current studies have focused on the humoral immune response elicited by this vaccine. This focus reflects the findings from clinical trials that protection from *P. falciparum* infection closely correlates with the concentration of specific antibody^12,56^. A focus on the humoral immune response also allowed us to correlate the concentrations of antibody attained in macaques with protection observed in the mouse model system.

This challenge model employs large inocula of sporozoites (5 x10^3^) compared to the challenge typically encountered following a mosquito bite (fewer than 200 sporozoites). Because this challenge ensures a large sporozoite load in the liver, it enables comparisons between different vaccination approaches by providing results that evaluate the relative efficacy of different approaches in reducing liver sporozoite load, viewed as a continuous variable. Challenge models employing lower inocula and evaluating outcomes such as time to patency or prevention or lack of prevention of bloodstream infection would be less amenable to the quantitative comparisons we are making in the current study. Because of differences in the challenge inocula used in liver sporozoite load vs. sterilizing protection studies, it is difficult to extrapolate in a quantitatively meaningful manner how results in one model system would translate to the other. However, in previous studies of sterilizing protection in mice with the MCSP platform, we have demonstrated that this vaccine platform can elicit complete or near complete protection against development of bloodstream infection after mosquito bite challenge ^16^.

The transgenic *P. berghei* sporozoite used in our mouse challenge studies carries only the central repeat region of the *P. falciparum* CSP, although our vaccine includes almost the entire protein, as described in the Methods. It is possible that even greater reduction of sporozoite copy number would be observed if the target parasite carried other antigens to which antibody might have been elicited by our vaccine. However, the central repeat region of this parasite protein is highly conserved and thought to be the dominant target of protective antibodies ^57,58^. It would be the most appropriate target to evaluate for immunogenicity and ability to reduce sporozoite infection.

These studies indicate that this vaccine formulation, which employs clinically approved adjuvants and induces no antibodies to the autologous host-derived iDC targeting component, can elicit in infant macaques high concentrations of specific antibody 18 weeks after the final of two immunizations. Those sustained antibody concentrations are capable of inducing at least a 70% reduction in liver sporozoite load in a rigorous mouse challenge model system. One weakness of this initial immunogenicity study in macaques is that it did not include control recipients of CSP alone to specifically confirm that the chemokine provided the same enhanced immunogenicity that has been consistently observed in our mouse studies. Further studies, to include that control and based on promising immunization protocols for inducing sustained immune protection ^13,14,15^, will address the potential of this vaccine construct for providing long term protection in the group that suffers the most severe consequences of malaria infection.

## Supporting information

Figure S1. Polyacrylamide gel electrophoresis analysis of purified vaccine constructs. Protein samples were separated by 12% SDS- PAGE for visualizat

## Acknowledgements

These studies were supported by a grant from the Johns Hopkins Malaria Research Institute and the Bloomberg Philanthropies.

## Author contributions

KL performed all of the experiments, analyzed the data, contributed to the interpretation of results and to the writing of the manuscript. JG assisted in the performance of the experiments and in the analysis of the data. FZ assisted in the planning of experiments, the interpretation of results, and the writing of the manuscript, RBM conceived of the experiments, and was involved in the planning and analysis of experiments, as well as the writing of the manuscript.

## Competing Interests statement

RBM and JG have equity interests in a company involved in the development of a malaria vaccine. KL and FZ declare no competing interests.

## Data set availability

The datasets generated during and/or analysed during the current study are available from the corresponding author on reasonable request.

