## Supplementary figures and images for "A Chemokine-Fusion Vaccine Targeting Immature Dendritic Cells Elicits Elevated Antibody Responses to Malaria Sporozoites in Infant Macaques"

### Figure S1. Polyacrylamide gel electrophoresis analysis of purified vaccine constructs. Protein samples were separated by 12% SDS- PAGE for visualizat

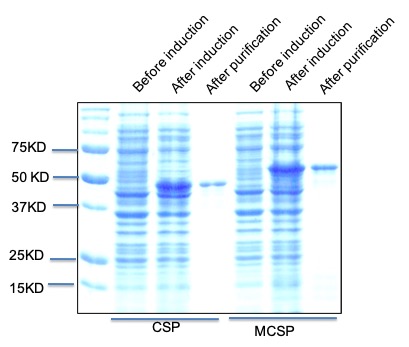
